# Cardiac performance and heat shock response variation related to shell colour morphs in the mudflat snail *Batillaria attramentaria*

**DOI:** 10.1101/2022.06.25.492793

**Authors:** Guodong Han, Yinghui Du, Lina Du, Furui Qu, Zhenjun Zhao

## Abstract

Physiological and transcriptomic response to thermal stress were investigated in the mudflat snail *Batillaria attramentaria* of different shell colour morphs. Despite no difference in upper lethal temperatures among shell colour morphs, their Arrhenius breakpoint temperatures (ABT) for cardiac thermal performance differed significantly, and the ABT of snails with D type morph was higher than snails with UL type morph. The transcriptomic analysis showed that D type snails exhibit higher levels of four HSPs than UL type snails at control temperature. Unfolded protein response was activated in UL type snails but not in D type snails under moderate thermal stress. And there were 11 HSPs increased in UL type snails in contrast to 1 HSP in D type snails, suggesting a “preparative defense” strategy of heat shock response in D type snails under moderate thermal stress. When exposed to sublethal temperature, eight molecular chaperones were uniquely upregulated in D type snails, suggesting these genes may allow for D type snails improved their cardiac thermal tolerance. The “preparative defense” strategies and higher ABT of cardiac thermal performance may adapt the dark shell snails to the rapid and stronger thermal stress in the field. Our results suggested that physiological selection imposed by moderate and sublethal thermal stress, instead of upper lethal limits, may be a driving force shaping shell colour frequency in the mudflat snail *B. attramentaria*.

## Introduction

Animal coloration is one of the most conspicuous phenotypic traits in natural populations and has important implications for adaptation (Poelstra *et al*. 2015, Smith *et al*. 2016). Both intertidal and terrestrial gastropods exhibit remarkable variation in the shell colour within and among populations, which is termed shell color polymorphism. Shell colour polymorphism serves taxonomists as characters that can be used to recognise and distinguish species, however their function for the gastropods is sometimes less clear and has been the focus of many ecological and evolutionary studies (reviewed in Williams, 2017). In many cases, shell colours of gastropods are frequently associated with environmental stresses, such as temperature (Etter 1988; Harris & Jones 1995; Miura *et al*. 2007; Phifer-Rixey *et al*. 2008), desiccation (Etter 1988) and salinity (Sokolova & Berger 2000). Additionally, long term studies have shown that variability of climatic selection have driven the change of shell colour frequency within and among populations (Ożgo & Schilthuizen 2012; Schilthuizen 2013). These works suggest that shell colour may affect fitness in gastropods and highlight the importance to understand the selective mechanisms for the maintenance of shell colour polymorphism, notably in the context of climate change.

Temperature affects all physiological and biochemical processes, translating into effects on metabolic processes, fitness and ecological dynamics (Angilletta 2009; Somero *et al*. 2017). Intertidal organisms frequently encounter extreme thermal stress during areal emersion, and solar radiation is usually the dominant component of the surface energy balance during low tide (Helmuth & Hofmann 2001; Seuront & Ng 2016), causing mortality in summer (Chan *et al*. 2006). Shell colour is known to affect body temperature and survivorship in intertidal gastropods. For example, when exposed to sunlight over a range of ecological relevant temperatures, the intertidal snail *Nucella lapillus* with brown morph suffered much greater mortality (Etter 1988). However, when snails were instead placed in a drying oven, the survivorship curves of brown and white morphs were quite similar. Similarly, shell colour was found to be a significant predictor of survivorship in the flat periwinkle *Littorina obtusata* when exposed to solar radiation, and snails with dark shells exhibited greater mortality relative to snails with light-coloured shells (Schmidt *et al*. 2007; Phifer-Rixey *et al*. 2008). In the manipulative trial in which snails were painted, original colour had no detectable effect (Phifer-Rixey *et al*. 2008). These results suggested that upper lethal temperatures were not associated with shell colour in these intertidal snails. Therefore, tolerance to extreme temperature may offer poor explanations for the shell colour frequency within and among populations.

Studies of intertidal gastropods have demonstrated that body temperatures may be influenced by shell colour, and individuals with dark shell morphs are more rapidly heated by solar radiation, and can reach higher body temperature than light shell morphs (Cook & Freeman 1986; Phifer-Rixey *et al*. 2008; Miller & Denny 2011). Organisms subjected to low levels of environmental stress when the conditions moderately deviate from the optima (Sokolova *et al*. 2012). Consequently, individuals with dark shell morphs may suffer moderate thermal stress more frequently than light shell morphs. At the organism level, heart rate (HR) increases with body temperature until the Arrhenius breakpoint temperature (ABT) is reached, after which HR decreases rapidly. For intertidal molluscs, ABT is not acutely lethal, but does reflect cumulative damage to the cell during the heating process (Han *et al*. 2013, 2017), and is used as a proxy for sublethal (but stress) temperature (Tagliarolo & McQuaid 2015; Dong *et al*. 2022). The differences in ABT reflect their distributions both in a large-scale temperature gradient (Tagliarolo & McQuaid 2015) and microhabitats within site (Li *et al*. 2021), and highlight the critical importance of sublethal effects differences in physiology. At the cellular level, moderate stress induces a set of transcriptomic responses that include the repair of DNA and protein damage, cell cycle arrest or apoptosis, the removal of cellular and molecular debris generated by stress, and an overall transition from a state of cellular growth to one of cellular repairs (Sokolova *et al*. 2012). Transcriptomic studies thus are providing insights into shell colour related differences under moderate thermal stress. The organism and cellular processes under moderate thermal stress are energetically costly and may divert energy flux and metabolic power from fitness-related functions and may be a driving force shaping shell colour frequency in gastropods.

The mudflat snail *Batillaria attramentaria* (previous referred to as *B. cumingi* in literature) is widely distributed along the Northwestern Pacific coast (Ozawa *et al*. 2009; Ho *et al*. 2015). The snail is a dominant species in the tidal flats, and plays an important role in ecosystem, because of its impact on ecosystem carbon flows (Kawasaki *et al*. 2019). The snail *B. attramentaria* exhibits remarkable variation in the shell colour within and among populations (Miura *et al*. 2007). A dark unbanded shell (D type morph) and a shell with a white line on the upper side of each whorl (UL type morph) were found in *B. attramentaria* from China coast. A previous study has found that geographical variations in shell colour polymorphisms in *B. attramentaria* were significantly correlated with the temperature of the locality of the population, suggesting thermal selection was one of the significant factors maintaining shell colour polymorphism (Miura *et al*. 2007). In the present study, we hypothesized that response to moderate and sublethal thermal stress may differ between snails with different shell colours. To test this hypothesis, we investigated the effects of acute changes in temperature on heart rate and gene expression level of two type morph individuals. We also determined the effect of acute high temperature exposure on mortality to test whether the upper thermal limits differ between snails with D and UL type morph. Our results may help to understand the selective mechanisms for the maintenance of shell colour polymorphism in gastropods.

## Materials and methods

### Samples collection

Samples were collected from a muddy shore located at Yangmadao Island, Shandong province, China (37°27’N, 121°36’E) at daytime low tides. To collect *B. attramentaria* snails randomly, an 18 cm × 18 cm quadrat was thrown in several directions and all snails within the quadrat were collected. This operation was repeated three to four times in order to examine the colour variation of the shell. To investigate the seasonal fluctuations of frequencies of shell colour patterns, samples collections were repeatedly conducted in summer (July 2021) and winter (January 2022). Differences in frequencies of shell colour patterns were analyzed using Chi-square test in R.

### Survival analyses

The effect of acute high temperature exposure on mortality was determined using D and UL type morph snails collected in summer (July 2021) and winter (January 2022). Snails were held in air in test tubes (25 mm diameter) placed in a water bath (Grant TXF 200, Grant, UK) and heated at a rate of 6°C·h^-1^. Survival exposure to 43, 46, 49, 50, 51, 52, 53°C was assessed from three groups of 4-5 snails. Following heat stress, the test tubes containing the snails were immersed in flow-through seawater at ambient temperature during a 3 days recovery period. Individuals that did not exhibit an opercular reflex (rapid and complete withdrawal into shell) upon stimulation by a sharp probe on the foot were scored as dead. The median lethal temperature (LT50) at 3 days after heat shock were calculated with Logistic analysis. The complete set of survival time data was analyzed with a Cox proportional hazard regression model using *survival* package in R.

### Thermal sensitivity of heart rate

The cardiac performance, which describes the relationship between body temperature and heartbeat, was determined using D and UL type morph snails collected in summer (July 2021). Heartbeats were measured using a non-invasive method (Dong *et al*. 2021). Snails were attached to the bottom of test tubes and treated by heating the bottom of test tubes in a Grant water bath. Experimental temperatures were increased from 28 °C at a rate of 6 °C·h^-1^ in air until a temperature was reached where the heart rate fell to zero. To measure the snail’s body temperature, a small hole (0.8 mm diameter) was drilled in the shell at a position above the heart, and a thermal couple was inserted inside the shell. The heartbeat was detected by means of an infrared sensor fixed to the shell at a position above the heart (upper left to the aperture). Variations in the light-dependent current produced by the heartbeat were amplified, filtered and recorded using an infrared signal amplifier (AMP03, Newshift, Portugal) and Powerlab AD converter (8/30, ADInstruments, Australia). Data were viewed and analyzed using LabChart v7 (ADInstruments, Australia). Arrhenius Breakpoint temperature (ABT) for cardiac performance, the temperature at which the HR (beats per minute) decreases sharply with progressive heating, is determined using a regression analysis method that generates the best fit line on either side of a putative break point for the relationship of ln-transformed HR against reciprocal value of absolute temperature. ABT was calculated using *segmented* package (Muggeo 2008) in R. Comparisons of ABT between individuals with D type and UL type morph were performed using t test in R.

### RNA sequencing

To analyze the transcriptomic response to thermal stress, snails of each morph were randomly selected and heated at a rate of 6°C·h^-1^ in air from 28°C to 36°C (moderate thermal stress) or 46°C (sublethal thermal stress). After heating, three snails of each morph were dissected and foot muscle were immediately frozen in liquid nitrogen. Tissue samples were sent to Novogene (Tianjin, China) and then sequenced on an Illumina NovaSeq platform. All clean reads from 18 transcriptomes of snail were aligned to the *B. attramentaria* genome (GenBank assembly accession: GCA_018292915.1) using *STAR* v2.7.9a. The mapped reads of each sample were transformed into counts using *HTSeq* v1.99.2. The differential expression analyses were conducted between the control (28°C) and the treatments (36°C or 46°C) for each morph. The absolute value of log2fold change ≥ 1 and the *p*-adjust value <0.01 were set as the thresholds to screen out the differential expression genes (DEGs). The differential expression analysis was also performed between the D type morph and the UL type morph at 28°C. Gene Ontology (GO) and KEGG enrichment analyses were applied using *clusterProfiler* v4.2.2 in R to determine the significantly enriched GO terms and KEGG pathways of DEGs. Enriched GO terms and KEGG pathways were screened out with adjusted *p*-value < 0.05.

## Results

### Abundance and frequencies of shell morph in the field

At the study site, the total abundance of *B. attramentaria* averaged 1672 ± 290m^-2^, of which D type morph constituted 82.8% and UL type morph constituted 17.2% in summer. Lots of empty shells were found along the coast in winter, and live snails tend to aggregate together. The total abundance of snails averaged 3138 ± 1394m^-2^, of which D type morph constituted 85.2% and UL type morph constituted 14.8% in winter. There is no difference of frequencies of shell morph between summer and winter (n = 675, χ^2^ = 0.218, *P* = 0.640).

### Survival following thermal stress

After 3 days recovery, the LT50 values of snails with D and UL type morph in summer were 50.288 ± 0.178°C (mean ± SE) and 49.921 ± 0.292°C, respectively (Fig. 1). In winter, the LT50 values of snails with D and UL type morph were 50.432 ± 0.301°C and 49.915 ± 0.386°C, respectively. The proportional hazard assumption was tested for each variable (temperature, shell morph, season and shell morph × season) of the fitted Cox model by correlating the status (alive or dead) with time (Table 1). Survival didn’t vary among shell morph (*P* = 0.5), season (*P* = 0.4) or shell morph × season (*P* = 0.8), except for temperatures (*P* < 0.001).

**Table 1.**
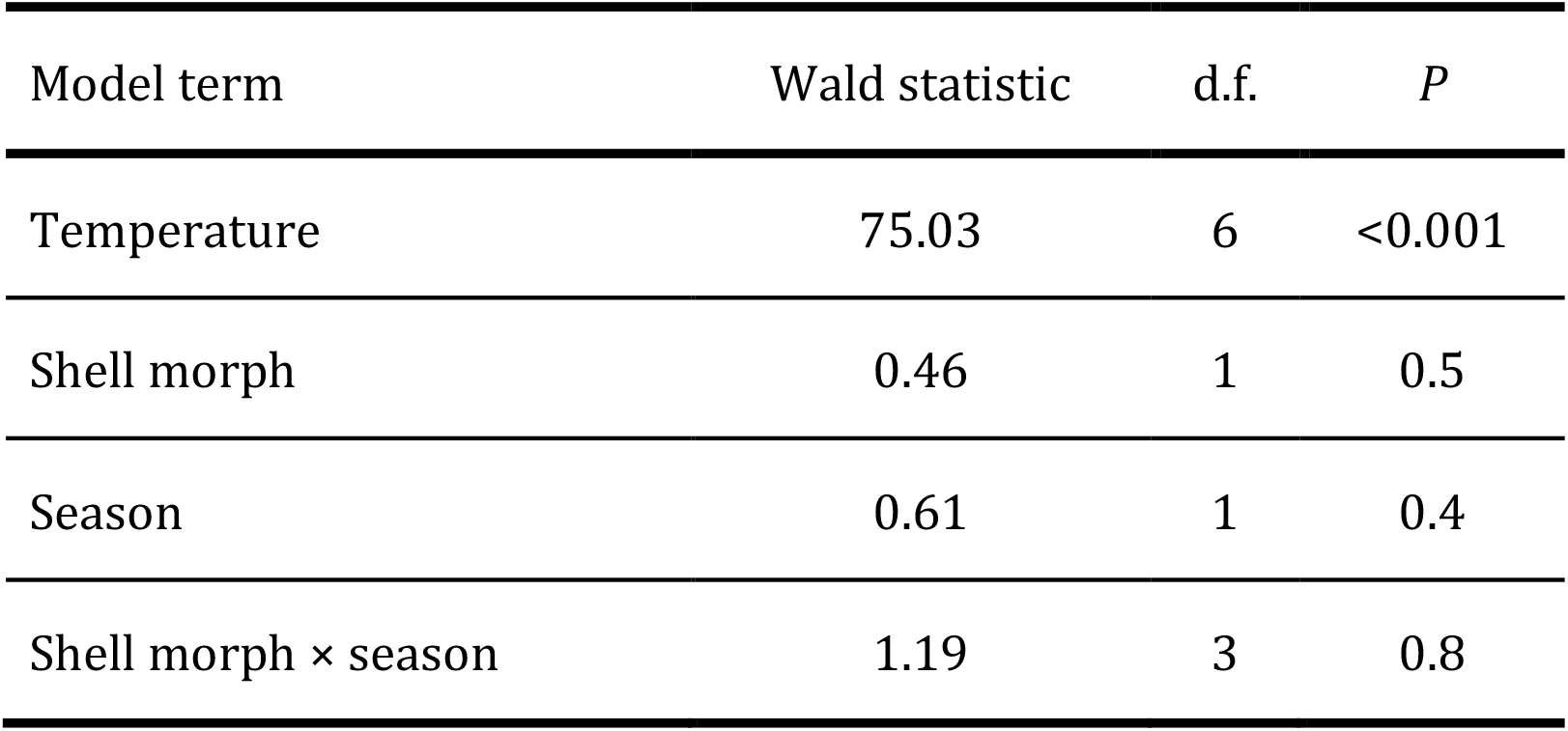
Statistical summary of the Cox proportional hazard model results for survival of snails after exposure to elevated body temperature

| Model term | Wald statistic | d.f. | *P* |
| --- | --- | --- | --- |
| Temperature | 75.03 | 6 | <0.001 |
| Shell morph | 0.46 | 1 | 0.5 |
| Season | 0.61 | 1 | 0.4 |
| Shell morph × season | 1.19 | 3 | 0.8 |

**Figure 1.**
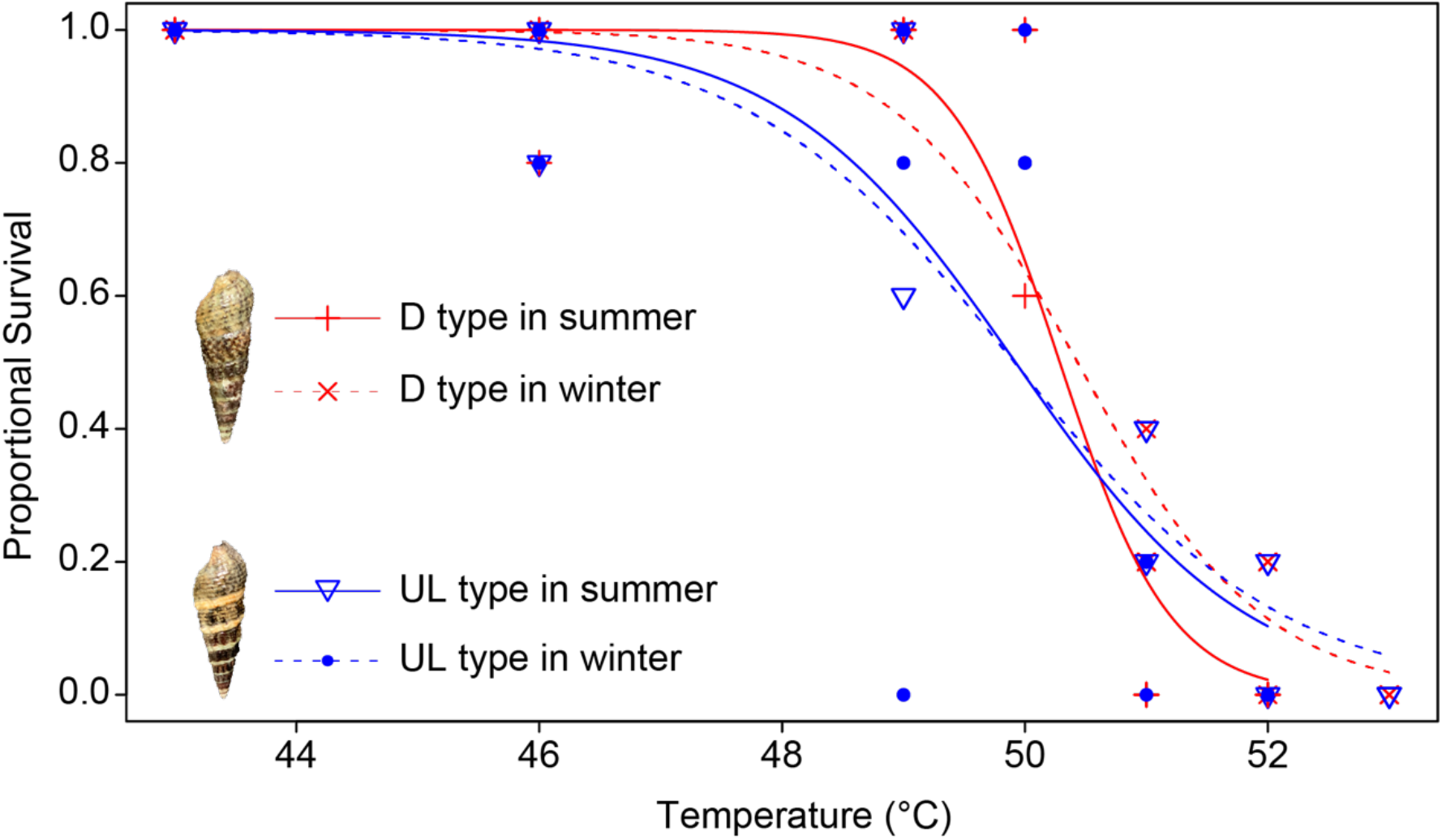
Survival rate of individuals of *B. attramentaria* after exposure to elevated body temperature. The curves were generated using Logistic regression.

### Cardiac thermal performance

Under constant heating, the patterns of heart rate were similar between two morphs when the temperature was below 40°C (Fig. 2). There was a flattening of the curve for both D and UL type morph. Cardiac thermal performances were different between two morphs when the temperature is above 40°C, especially when the temperature reached to ABT. The Means (±SD) of ABT for individuals with D and UL type morph were 45.627 ± 0.561 and 44.790 ± 0.894°C, respectively (Fig. 3a). The ABT of individuals with D type morph was higher than individuals with UL type morph (two sample t-test, *t* = 2.631, df = 20, *P*-value = 0.016). The means (±SD) of maximum heart rate (MHR) for each morph were: 136.727 ± 17.281 (D type) and 147.727 ± 16.900 beats min^-1^ (UL type, Fig. 3b). There was no significant difference in MHR between two morphs (two sample t-test, *t* = -1.509, df = 20, *P* = 0.147).

**Figure 2.**
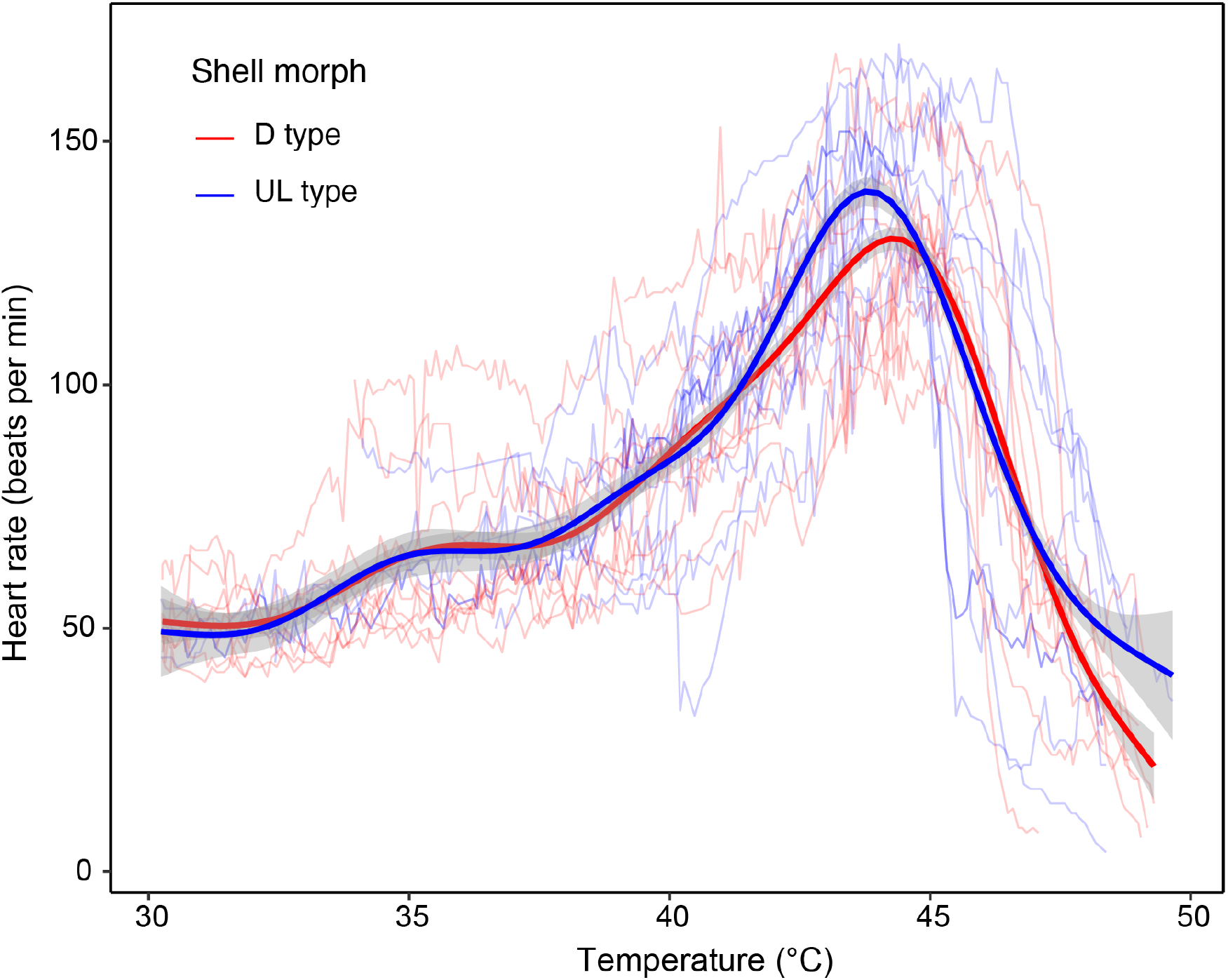
Cardiac thermal performance of individuals of B. attramentaria over an acute warming ramp. Each thin transparent line represents a cardiac performance of an individual with D (red) or UL (blue) type morph over different temperatures. The thick red and blue curve is fitted from GAM model for individuals with D (n=11) and UL (n=11) type morph, respectively, and grey regions are 95% confidence intervals.

**Figure 3.**
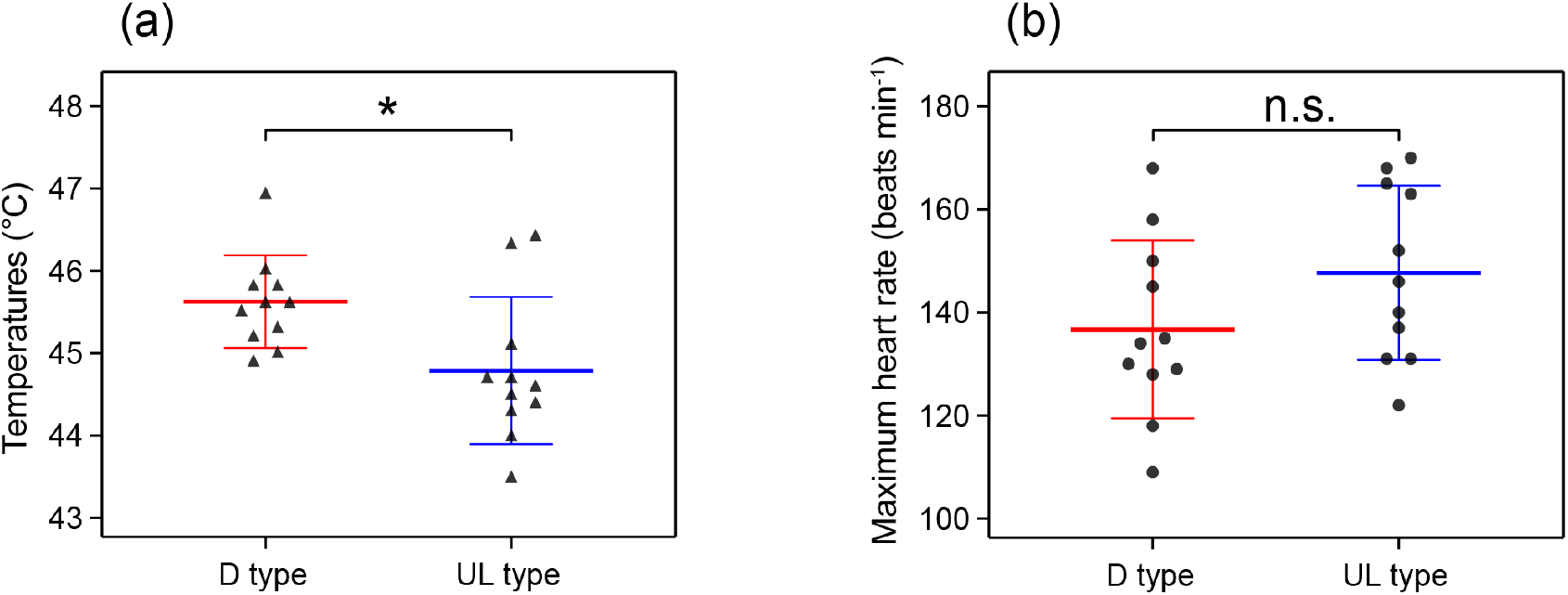
The mean ABT and maximum heart rate of B. attramentaria (mean ± SD). (a) Each triangle represents individual ABT (n=11 for both D and UL type morph). * indicates significant differences between morphs (*P*=0.016). (b) Each point represents individual maximum heart rate (n=11 for both D and UL type). There is no significant different between morphs (*P*=0.147).

### DEGs under temperature stresses

A total of 121.70 Gb clean bases were obtained from 18 transcriptomes of *B. attramentaria*. The average mapping rates for D and UL type morph individuals were 85.39% and 84.09%, respectively. At control temperature (28°C), the expression levels of 37 genes were higher in D type than UL type morph individuals, while 68 genes were lower in D type than UL type morph individuals (Fig. 4). In response to 36°C, there were 84 DEGs (46 up-regulated and 38 down-regulated) in D type morph individuals in contrast to 37 DEGs (23 up-regulated and 14 down-regulated) in UL type morph individuals. When the temperature increased to 46°C, the number of DEGs were 1054 (641 up-regulated and 413 down-regulated) and 669 (475 up-regulated and 194 down-regulated) for D and UL morph individuals, respectively.

**Figure 4.**
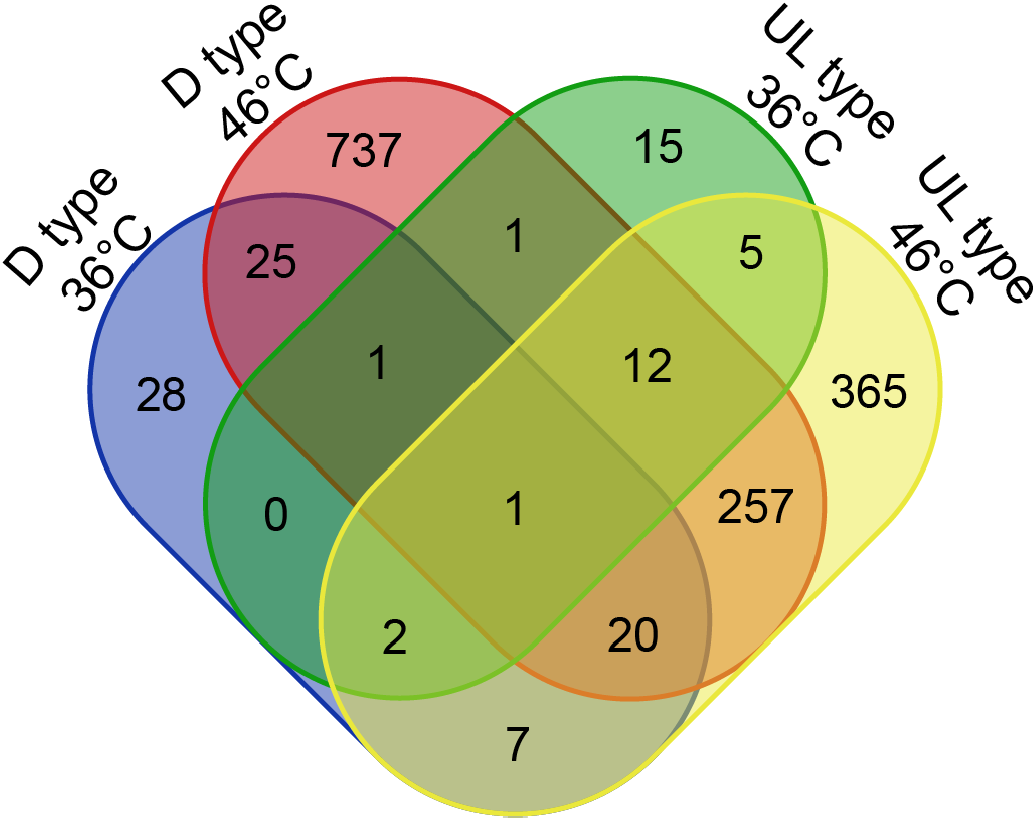
Venn diagram comparing the number of DEGs in response to heat stress at 36°C and 46°C in D and UL type morph individuals.

### Functional analysis of DEGs

GO annotations for the DEGs were performed under heat stress (Fig. 5a). In response to 36°C, 38 DEGs and 21 DEGs were annotated in the GO database in D type and UL type morph individuals, respectively. The terms “primary amine oxidase activity”, “amine metabolic process”, “quinone binding” and “myosin complex” were enriched in D type morph individuals. The term “ATP hydrolysis activity” turned out to be the most enriched GO term in UL type morph individuals, followed by “unfolded protein binding” and “protein folding”. When the temperature increased to 46°C, terms “ATP hydrolysis activity”, “unfolded protein binding” and “protein folding” were enriched in both D and UL type morph individuals. GO terms “DNA integration” and “calcium ion binding” were uniquely enriched in D type morph individuals, and “ endopeptidase inhibitor activity” and “signaling receptor activity” uniquely enriched in UL type morph individuals.

**Figure 5.**
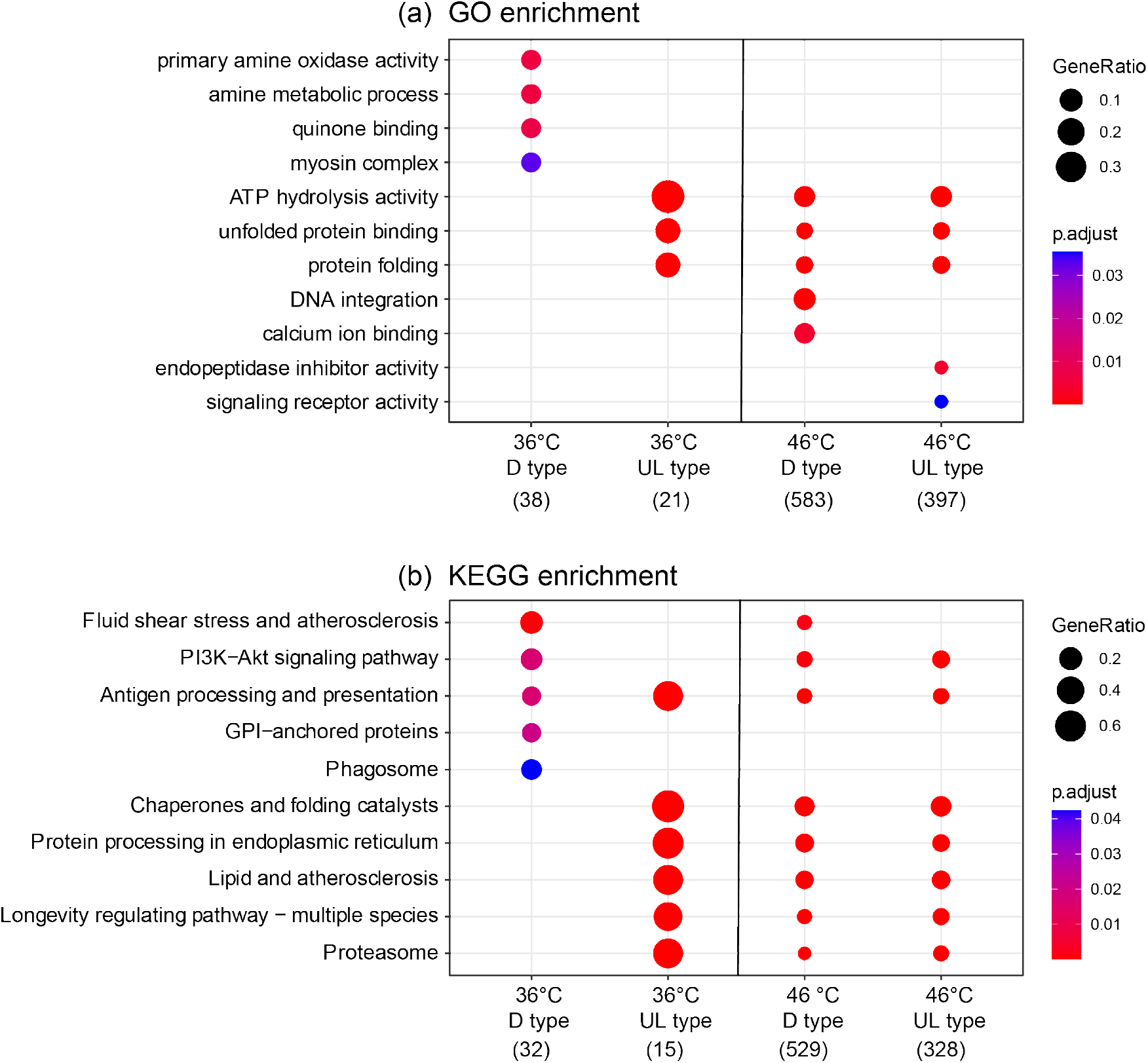
DEGs enriched in the top GO terms (a) and KEGG pathways (b). The number in brackets represents the total number of DEGs annotated in functional classes (GO terms or KEGG pathways) in response to heat stress at 36°C and 46°C in D and UL type morph individuals. The dot size represents the gene ratio between the number of DEGs enriched in each category and the total number of DEGs annotated in functional classes. The *p* adjust values were shown in colors, the red color being more significant than the blue color.

In response to 36°C, KEGG annotations revealed that 32 DEGs and 15 DEGs were mapped to pathways in D type and UL type morph individuals, respectively (Fig. 5b). Among these pathways, “Fluid shear stress and atherosclerosis” was the most significantly enriched pathway in D type morph individuals, containing 6 DEGs. And “Chaperones and folding catalysts” was the most significantly enriched pathway in UL type morph individuals, containing 10 DEGs. When the temperature increased to 46°C, KEGG annotations revealed that 529 DEGs and 328 DEGs were mapped to pathways in D type and UL type morph individuals, respectively. The pathway “Chaperones and folding catalysts” was the most significantly enriched pathway in both D and UL type morph individuals.

### Heat shock protein genes

The expression levels of 4 heat shock protein (HSP) genes, which were annotated as HSP70B2 and DNAJB1, were higher in D type morph individuals than UL type at control temperature (Fig. 5a). There was only 1 HSP gene (HSP90A1) upregulated in D type morph individuals in response to 36°C in contrast to 11 HSP genes in UL type morph individuals. These 11 HSP genes were annotated as DNAJA1, HSP90A1, DNAJB1, HSP70B2, CRYAA, HSPIV and HSPA8. When the temperature increased to 46°C, a total of 35 HSP genes were upregulated in D and UL type morph individuals, of which 26 HSP genes were both upregulated in D and UL type morph individuals. There were 8 HSP genes uniquely upregulated in D type morph individuals in contrast to 1 unique HSP gene in UL type morph individuals.

## Discussion

Our study was designed to examine the correlation between shell colour and physiological and transcriptomic responses to thermal stress in an intertidal gastropod *B. attramentaria*. We asked whether responses to moderate and sublethal thermal stress were associated with shell colour polymorphism in gastropods. Using heart rate, gene expression levels and mortality as proxies, we showed that responses to moderate and sublethal thermal stress, instead of lethal thermal stress, seem to be associated with shell colour morph in *B. attramentaria*.

The D type snails frequently appeared on the study site (contributing 82.8% of total colour morphs in summer), suggesting a fitness advantage for D type morphs on the study site. Differences in mortality rate of different morphs of polymorphic gastropods may be causally related to the variation in their shell colour (Köhler *et al*. 2021). However, the survivorship didn’t differ between D and UL type morph snails when exposed to acute heat stress in laboratory. Our results were consistent with the previous studies in intertidal snails *Nucella lapillus* (Etter 1988) and *Littorina obtusata* (Phifer-Rixey *et al*. 2008), which showed that upper lethal temperatures were not associated with shell colour. These results indicated that physiological selection imposed by extreme temperature conditions may not be a driving force shaping shell colour frequency of *B. attramentaria*.

The present study showed that D type snails exhibited higher ABT than UL type snails. Although the relationship between shell colour patterns and body temperature of *B. attramentaria* snails wasn’t examined in the present study, other studies of intertidal gastropods have demonstrated that individuals with dark shell morphs are more rapidly heated by solar radiation, and can reach higher body temperature than light shell morphs (Cook & Freeman 1986; Miller & Denny 2011; Köhler *et al*. 2021). Thus, snails with D type shell may suffer stronger thermal stress than snails with UL type shell in the field. Our findings are similar to previous research in that they demonstrated that ABT of intertidal molluscs inhabiting hot conditions was higher than molluscs inhabiting benign conditions within a population (Moyen *et al*. 2019; Li *et al*. 2021). ABT is not acutely lethal, but does reflect cumulative damage to the cells that is initiated during earlier stages of heating and gradually builds up to a level that causes heart dysfunction at the critical temperature (Han *et al*. 2013; Han *et al*. 2017). The ABT of cardiac function, while an index of organ-level dysfunction, thus can serves as an indicator that sufficient thermal damage of cellular structures has occurred to render the heart suboptimal in its performance (Moyen *et al*. 2019). Higher ABT indicated that D type snails may be better adapted to thermal conditions around sublethal temperature. The observed variability in cardiac thermal performance suggested that there are likely cellular and molecular changes allowing for the snails with D type shells improved cardiac thermal tolerance.

When exposure to moderate thermal stress (36°C), GO terms including “unfolded protein binding” and “protein folding” and KEGG pathways including “chaperones and folding catalysts” and “protein processing in endoplasmic reticulum” were significantly enriched in UL type snails but not in D type snails. These results suggested that the unfolded protein response was activated in only UL type snails in response to moderate thermal stress. To ascertain fidelity in protein folding, cells regulate the protein-folding capacity in the endoplasmic reticulum according to need (Hetz 2012). The endoplasmic reticulum responds to the burden of unfolded proteins in its lumen by activating intracellular signal transduction pathways (Walter & Ron 2011). The lack of unfolded protein responses in D type snails suggested that when these snails are exposed to moderate thermal stress, they do not mount a transcriptome-wide intracellular signal transduction pathways (which may be very energy costly) and only induce genes essential to address immediate damage. The energy saving process may be a result of timing metabolic depression (Hui *et al*. 2020), which can allow intertidal gastropods depress resting metabolism in response to moderate thermal stress (Marshall *et al*. 2011; Chen *et al*. 2021). Additionally, the decrease observed for the gene phosphoenolpyruvate carboxykinase at 36°C was further evidence for a decrease in metabolism in response to moderate heat stress (Tomanek & Zuzow 2010). Therefore, these snails may benefit from temporal constraint on energy gain while experiencing high body temperature.

As observed in functional enrichment analyses, significant overexpression of several molecular chaperones was detected in response to thermal stress, which is in line with previous studies in intertidal gastropods (Sorte & Hofmann 2004; Wang *et al*. 2014; Gleason & Burton 2015; Han *et al*. 2017). We identified all differentially expressed HSPs and cofactors, and found distinct strategies of HSPs in *B. attramentaria* snails with different shell colours. At acclimation (control) temperature, the expression levels of four HSP70B2 paralogs and DNAJB1 (HSP40) were significantly higher in D type snails than UL type snails (Fig. 6a). Dong *et al*. (2008) found that high-intertidal congeners of *Lottia* employ a “preparative defense” strategy involving maintenance of high constitutive levels of Hsp70 in their cells as a mechanism for protection against periods of extreme and unpredictable heat stress. Our data suggested that *B. attramentaria* snails with D type shell may benefit from such “preparative defense” strategies in response to moderate stress. Several molecular chaperones, including five HSP70B2 paralogs, HSP70IV, CRYAA, HSPA8, DNAJA1 and DNAJB1, were not differentially expressed in D type snails under moderate thermal stress compared to UL type snails. Our data generally agreed with previous results that found less thermal tolerant gastropods under benign conditions show higher HSP protein expression following heat stress than congeners under warmer conditions (Tomanek & Somero 1999, 2000; Tomanek 2010). When exposure to 46°C, eight molecular chaperones, including DNAJC16, CRYAB, HSPA14B, DNAJC3, two HSPA5 paralogs, DNAJC10 and HSP90B1, were uniquely upregulated in D type snails. This suggested that evolution of elevated expression of these genes under extreme thermal stress adapts the D type snails to the rapid and stronger thermal stress.

**Figure 6.**
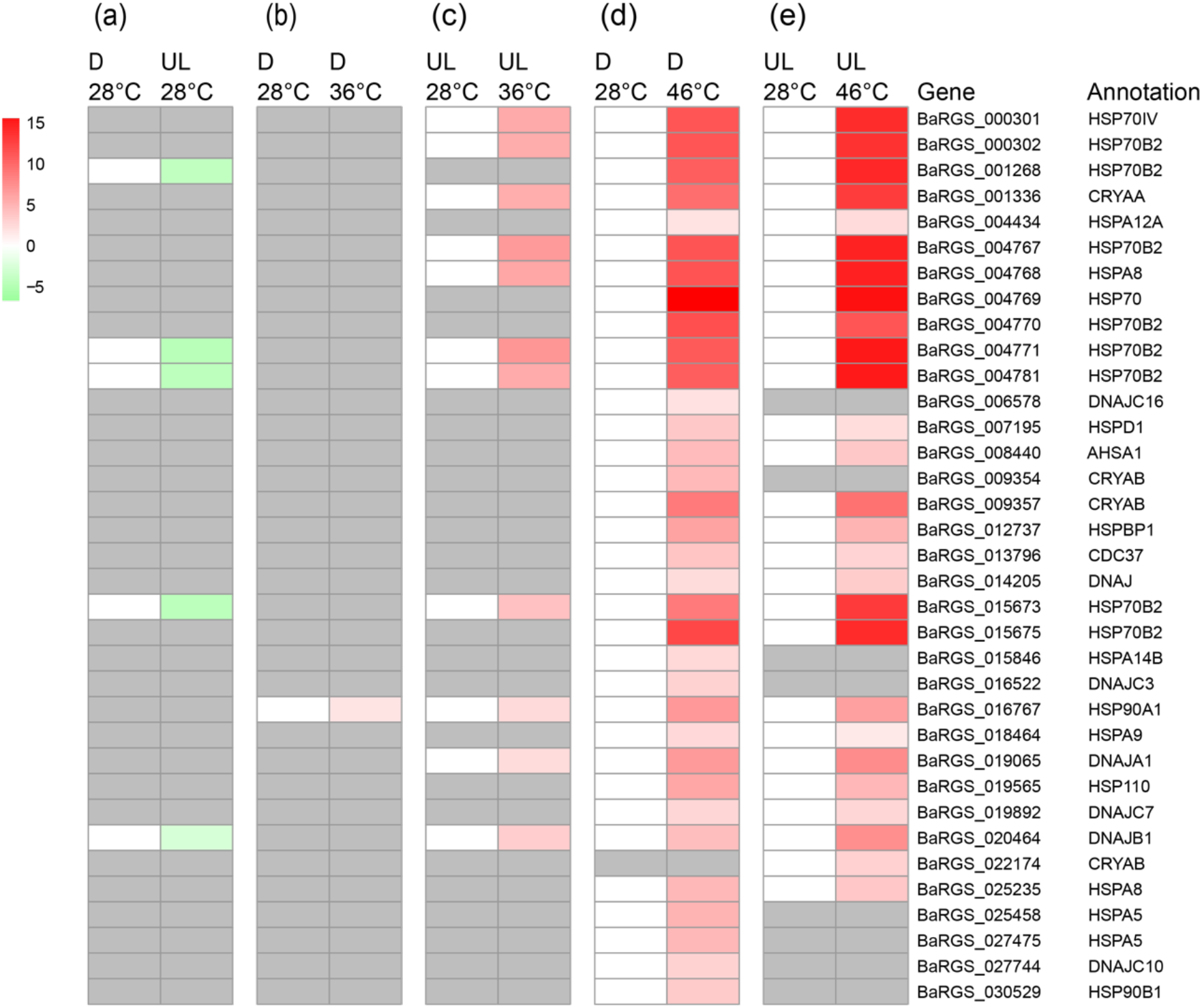
Heatmap of differentially expressed heat shock genes. In each group, the differential expression analyses were conducted between the control (first column) and treatment (second column). Each differentially expressed gene is represented by a single row, and the log2 fold change values are shown in colors. Red corresponds to up-regulated gene product, and green corresponds to down-regulated gene product. Grey represents no significant difference.

The patterns of shell colour polymorphism can be affected by a number of selective processes such as visual selection (Heller 1975), sexual selection (Rolán-Alvarez *et al*. 2012), thermal regime (Schilthuizen 2013) and balancing selection (Johannesson & Butlin 2017). A previous study has found that thermal regime profoundly contributes to maintaining the shell colour polymorphisms in *B. attramentaria* (Miura *et al*. 2007). Our study provides an example of the potential for physiological selection imposed by moderate and sublethal thermal stress to shape shell colour polymorphism. The potential impact of moderate temperature as a selective force is especially significant. Moderate thermal stress is not immediately lethal, but does divert energy flux from fitness-related functions such as reproduction and growth towards maintenance and repair (Sokolova *et al*. 2012; Han *et al*. 2013). *B. attramentaria* is adapted to a broad range of temperatures throughout the intertidal regions of the Northwestern Pacific (Ozawa *et al*. 2009; Ho *et al*. 2015). The results of our study suggested that global warming has the potential to substantially change the distribution of shell colour morph frequencies in *B. attramentaria* along the Northwestern Pacific coast.

## Conclusion

Our study provides an example of the potential for thermal selection for the maintenance of shell colour polymorphism in gastropods. The mortality didn’t differ in the snails with different shell coloration in response to lethal temperature. However, transcriptomic analysis suggested that unfolded protein response was anly activated in light shell snails under thermal stress. Snails with dark shell exhibit high levels of specific HSPs at control temperature, and may employ a “preparative defense” strategy. The mean ABT was higher in dark shell snails than light shell snails, indicating dark shell snails can maintain cardiac performance at higher temperature. When exposed to sublethal temperature, eight molecular chaperones were uniquely upregulated in D type snails, indicating these genes may allow for the snails with D type shells improved cardiac thermal tolerance. Our results suggested that physiological selection imposed by moderate and sublethal thermal stress, instead of upper lethal limits, may be a driving force shaping shell colour frequency in the mudflat snail *B. attramentaria*.

